# Temporal Dynamics of Apoptosis-Induced Proliferation in Pupal Wing Development: Implications for Regenerative Ability

**DOI:** 10.1101/2023.07.10.548326

**Authors:** Sara Ahmed-de-Prado, Carlos Estella, Antonio Baonza

## Abstract

The ability of animals to regenerate damaged tissue is a complex process that involves various cellular mechanisms. As animals age, they lose their regenerative abilities, making it essential to understand the underlying mechanisms that limit regenerative ability during aging. *Drosophila melanogaster* wing imaginal discs are epithelial structures that can regenerate after tissue injury. While significant research has focused on investigating regenerative responses during larval stages, particularly regarding the regulation and function of the JNK pathway, our comprehension of the regenerative potential of pupal wings and the underlying mechanisms contributing to the decline of regenerative responses remains limited. This study explores the temporal dynamics during pupal development of the proliferative response triggered by the induction of cell death, a typical regenerative response. Our results indicate that the apoptosis-induced proliferation response can be initiated as late as 30 hours after pupa formation (APF), when in normal circumstances cell proliferation ceases at around 20 hours APF. Furthermore, our data revealed that after 35 hours APF, cell death alone fails to induce further proliferation. Interestingly, the failure of reinitiating the cell cycle beyond this time point is not attributed to an incapacity to activate the JNK pathway. Instead, one of the constraining factors in the apoptotic-induced proliferation process during pupal development seems to be the activity level of ecdysone-responsive genes.

**Author Summary:** Animals have the remarkable ability to regenerate damaged tissues, but this regenerative potential diminishes with age. Understanding the mechanisms underlying age-related decline in regenerative abilities is crucial. Drosophila melanogaster wing imaginal discs provide a valuable model for studying tissue regeneration. While significant research has focused on regenerative responses during larval stages, our understanding of the regenerative potential and mechanisms in pupal wings remains limited.

In this study, we investigate the temporal dynamics of the proliferative response triggered by cell death during late during the development, in pupal development. Our findings reveal that the apoptosis-induced proliferation response can occur during pupal development, even after normal cell proliferation has ceased. However, at late stages of pupal development this response does not occur. We have found that, the inability to reinitiate the cell cycle beyond this time point is influenced by the activity of the hormone ecdysone and its-responsive genes.

These findings shed light on the dynamic processes involved in tissue regeneration during pupal development. This study expands our understanding of the complex interplay between cell death, proliferation, and gene activity during tissue regeneration, providing valuable insights for future research in regenerative biology.

## Introduction

Regeneration is a remarkable ability present across the animal kingdom that enables multicellular organisms to repair damaged tissues and maintain tissue homeostasis (Tanaka and Reddien, 2011;Brockes and Kumar, 2008). This intricate process involves various cellular mechanisms, including regenerative growth. As animals age, they progressively lose their regenerative abilities, including some salamanders with boundless regenerative capacities. Understanding the underlying mechanisms that limit regenerative ability during aging is a crucial developmental biology question.

During development, intrinsic cellular and physiological changes occur that limit the ability of signaling pathways to promote cellular plasticity and induce regenerative growth, which are crucial to support regeneration (McCusker and Gardiner, 2011; Seifert and Voss, 2013). For instance, differentiated cells lose the ability to re-enter the cell cycle, a process necessary for limb regeneration in vertebrates. Tumor suppressor proteins like the Retinoblastoma protein (Rb) likely play a crucial role in maintaining plastic cellular states and cell cycle re-entry. The levels of this factor vary during regeneration in salamanders and across developmental stages in mammals (Seifert and Voss, 2013). However, the loss of the ability to regenerate limb buds can be delayed experimentally in *Xenopus laevis*, indicating that the loss of regenerative ability during aging may also be postponed or reversed.

The imaginal wing disc of *Drosophila melanogaster* is a well-established model tissue for studying epithelial damage repair (Bergantinos et al., 2010b; Fox et al., 2020; Ahmed-de-Prado and Baonza, 2018; Worley and Hariharan, 2022). These sac-like structures have the remarkable ability to regenerate during larval stages but lose this ability at the end of the larval stage or during pupal development, which coincides with the cessation of cell proliferation (Smith-Bolton et al., 2009; Diaz-Garcia and Baonza, 2013; Buttitta et al., 2007).

During pupal development, cells exit the cell cycle as a result of a decrease in Cyclin E (CycE) and an increase in Rb factor activity (Buttitta et al., 2007). Rb represses the transcription of genes required for the G1-S transition, including *CycE*, by binding and inhibiting the transcription factor E2F1 (Du et al., 1996a; Du et al., 1996b). CycE, in turn, promotes the G1-S transition by phosphorylating and inhibiting Rb. Although cell cycle exit occurs around 24 hours APF, the positive feedback loop between CycE and E2F1 is maintained until around 30-35 hours APF. Therefore, over-expression of either CycE or E2F1 between 24-35 hours after pupa formation (APF) is sufficient to induce proliferation. However, after 30-35 hours, co-expression of *CycE* and *E2F1* is necessary to induce proliferation (Buttitta et al., 2007). The positive feedback between CycE and E2F1 ends around the onset of epigenetic shutdown of regulatory regions of key genes involved in cell cycle control, such as *CycE*, *E2F1*, and *string* (Buttitta et al., 2007). Recent research has revealed that ecdysone-responsive transcription factors regulate the temporal changes in chromatin accessibility during wing disc development (Uyehara et al., 2017).

The c-Jun N-terminal kinase (JNK) signalling pathway has emerged as a critical signal in the process of regeneration. Upon injury, the JNK signalling pathway is initiated at the wound site (Bosch et al., 2005). This pathway plays a key role in regulating several biological processes associated with regeneration (Bosch et al., 2005; Bergantinos et al., 2010a; Bogoyevitch et al., 2010) (Lee et al., 2005) (Chen, 2012). In studies of disc regeneration, inhibition of JNK function was found to negatively affect wound healing and reduce regenerative proliferation (Bosch et al., 2005; Mattila et al., 2005; Ramet et al., 2002). During regeneration, JNK signalling promotes the activation of several downstream pathways, including JAK/STAT and Wingless (Wg) (Harris et al., 2016; Katsuyama et al., 2015; Smith-Bolton et al., 2009; Pastor-Pareja et al., 2008).

Despite extensive research on regenerative responses during larval stages, our understanding of the regenerative abilities of pupal wings and the mechanisms involved in the decline of regenerative responses remains limited. It is unclear whether the signals activated by tissue damage during larval stages also operate during pupal development.

In this study, we investigated the proliferative response triggered by the induction of cell death during pupal development. Our findings indicate that apoptosis-induced proliferation (AiP) response can be triggered up to 30 hours APF. As cell proliferation normally ceases at 20-24 hours APF during normal development, these results suggest that cell death can extend the proliferative phase of cells in the wing discs during pupal development. We found that after 35 hours APF cell death is not sufficient to induce proliferation. Our data suggest that the inability to reactivate the cell cycle after this time point is not due to an inability to trigger the JNK pathway. Rather, one of the limiting factors in the apoptosis-induced proliferation process during pupal development appears to be the activity of ecdysone-responsive genes.

## Results

### Ionizing Irradiation (IR) can trigger an apoptosis-induced proliferation response up to 30-35 hrs APF

During the larval stage, the cells in the wing disc proliferate asynchronously. However, once the larva-to-pupa transition occurs (0 hours APF), the cells in the wing disc undergo a G2 phase arrest, which stops their proliferation. By 6 hours APF, the majority of cells in the pupal wing have entered this G2 phase arrest. Between 12-18 hours APF, a fraction of these cells resumes the cell cycle and undergo the last round of division before being arrested in G1. By 24 hours APF, approximately 95% of the wing disc cells are arrested in G1 phase and become quiescent before initiating terminal differentiation (Milán et al., 1996b; Buttitta et al., 2007; Schubiger and Palka, 1987; Guo et al., 2016).

We analyse the proliferative response of wing disc cells following the induction of apoptosis during pupal development. To induce cell death, we employed X-ray irradiation on pupae at three different time intervals: 0-4, 10-14, and 20-24 hours APF. Subsequently, we dissected the pupae 20 hours after the irradiation process (Fig. 1A). To assess the proliferative response, we conducted an analysis of the mitotic marker Phospho-Histone 3 (PH3) and determined the mitotic index of wing discs in the pupal stage (see methods).

**Figure 1.**
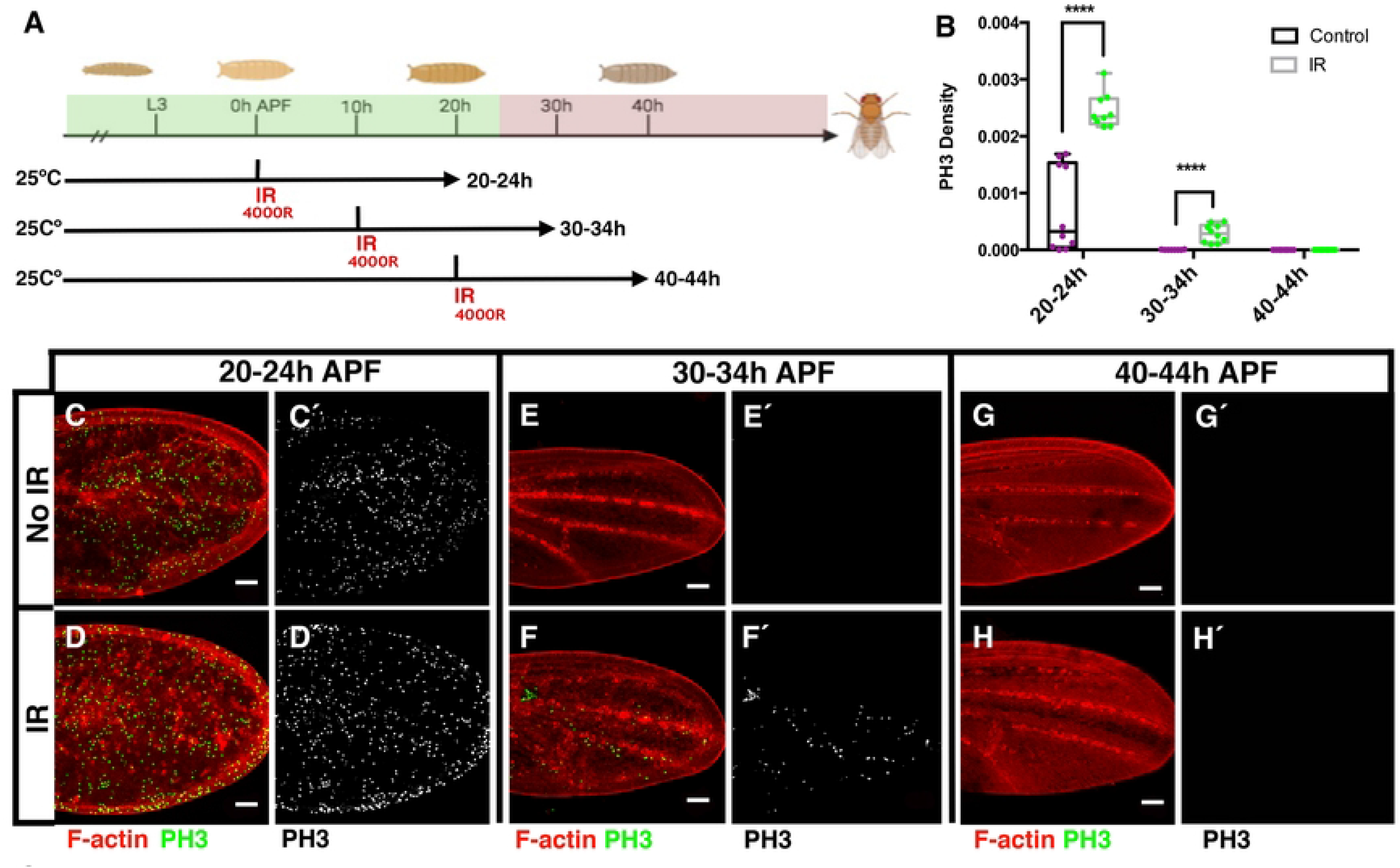
**Proliferative response in the pupal wing to X-ray irradiation at different time after puparium formation**. (A) Schematic diagram of the temperature shifts used in this experiment. The proliferative period of the pupal wing is represented by the green colour, while the period following cell cycle exit is indicated by the red colour. Here and in the rest of the figures the temperature treatment is described in the Materials and Methods section. (B) The graph shows the density of mitotic cells (PH3 positive) in control and irradiated pupae at different times APF. Error bars represent SEM. (C-C’, E-E’ and G-G’) Control pupal wing at 20-24hours (C-C’), 30-34 hours (E-E’) and 40- 45 hours (G-G’) after pupa formation APF. (D-D’, F-F’ and H-H’) Irradiated pupal wing at 20-24hours (D-D’), 30-34 hours (F-F’) and 40-45 hours (H-H’) after pupa formation APF (F-F’). Note that while there are no dividing cells in the control discs at 30-34 hours APF, cell proliferation is sustained in the irradiated discs at the same time point. The discs were stained with anti-PH3 antibody and Phalloidin. Statistical analysis was conducted using multiple comparation t-Test student (Mann-Whitney test) **** p < 0.0001. Control 20-24 hours n=10, 30-34 hours n=8 and 40-45 n=10 hours. Irradiated 20-24 hours n=9, 30-34 hours n =10 and 40-45 n=10 hours. White scale bar, 50μm

In wing discs examined at 20-24 hours (APF), we observed the presence of mitotic cells in both irradiated and non-irradiated pupae. However, the irradiated pupal wings had a higher mitotic index (Fig. 1B-D’). Surprisingly, we also observed a relatively high number of mitotic cells in irradiated pupal wings examined at 30-34 hours APF, even though there were no dividing cells in control discs at that time point (Fig. 1 E-F’). At 40-44 hours APF, neither the control nor the irradiated discs showed any mitotic cells (Fig. 1 G-H). These findings suggest that irradiation-induced apoptosis may prolong the proliferative period up to 30-34 hours APF.

IR-induced apoptosis depends on the proliferative status of the cells, with proliferating cells being more sensitive to irradiation than differentiating cells (Ruiz-Losada et al., 2022; Baonza et al., 2022). Therefore, the lack of a proliferative response at 40-44 hours APF upon IR might be due to cells no longer being sensitive to irradiation at this stage. To investigate this possibility, we examined radiation-induced apoptosis in pupal wing irradiated at 0-4 hours and 20-24 hours APF and examined 20 h later (20-24h and 40-44h APF, respectively). Our study revealed a notable increase in apoptosis in irradiated pupal wing cells at 20-24 hours APF (irradiated at 0-4 hours APF), compared to control discs, as indicated by the apoptotic marker Dcp-1 (see Fig. S1). However, we did not detect apoptotic cells in wing discs at 40-44 h APF (irradiated at 20-24 hours APF Fig. S1). These findings suggest that wing disc cells at 40 hours APF are insensitive to irradiation-induced apoptosis.

To further pinpoint the time at which pupal disc cells become insensitive to irradiation, we selected pupae at 2-hour intervals, starting at 20 hours APF (Fig. 2A). The selected pupae were immediately irradiated, and their apoptotic response was analyzed 4 hours later. Our findings revealed that the wings of pupae irradiated at 20-22 hours and examined 4 hours later showed a significantly higher number of apoptotic cells than control wings (Fig. 2 B-C). However, we did not observe an increase in the number of apoptotic cells in pupal wings of animals irradiated at 22-24h and 24-26 h APF (Fig 2 D-G). Therefore, these results suggest that the cells of pupal wing become radioresistant around 26 h APF concomitant to their definitive cell cycle arrest.

**Figure 2.**
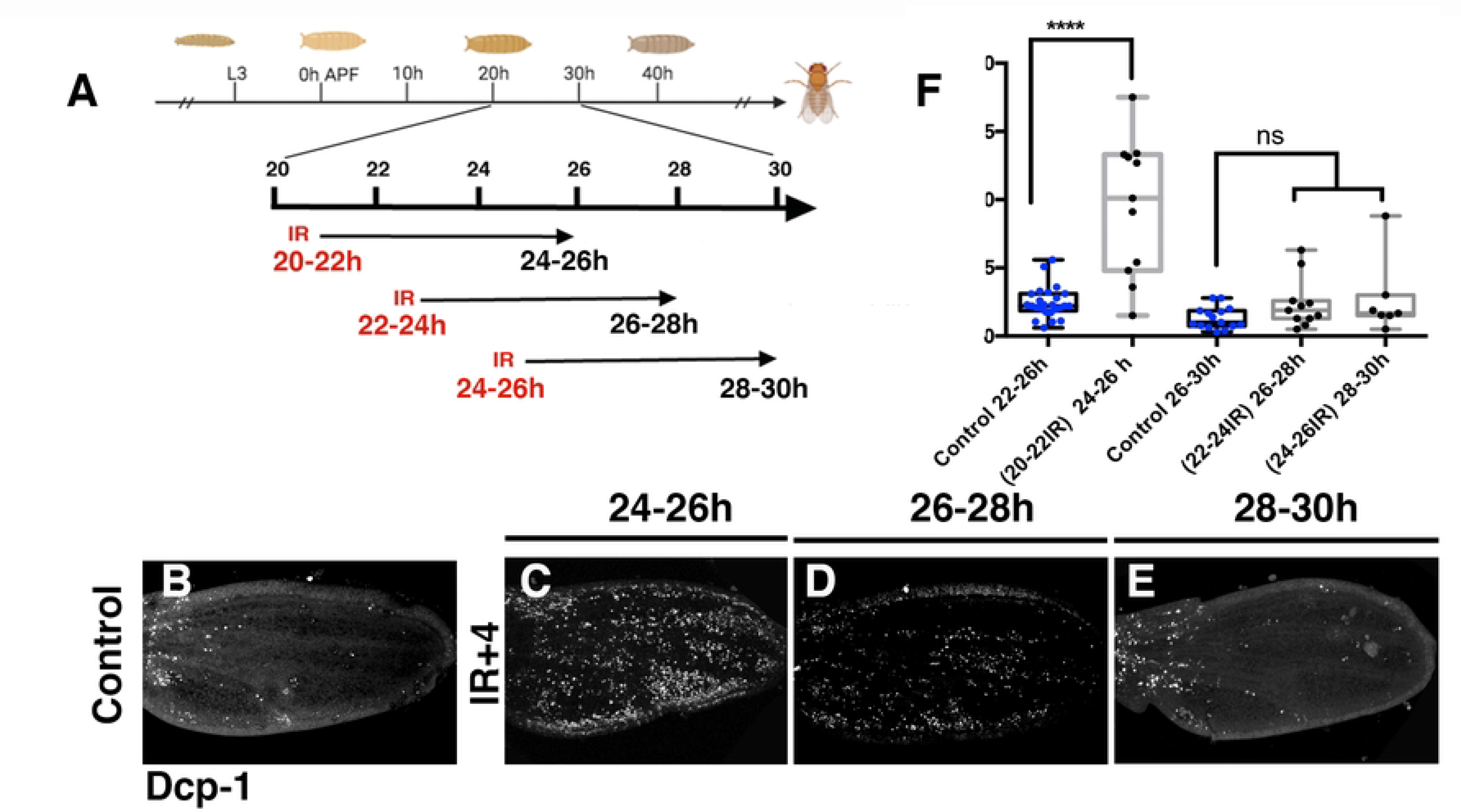
**Cell death induction by irradiation during pupal development**. (A) Schematic diagram of the irradiation times employed in this experiment. (B) Control pupal wing at 24-26 hours APF. (C, D, and E) Irradiated pupal wing at 24-26 hours (C), 26-28 hours (D), and 28-30 hours (E) APF. (F) The graph shows the density of apoptotic cells (Dcp-1 positive) in control and irradiated pupae at different times APF. Error bars represent SEM. The discs were stained with anti-Dcp-1 antibody. Statistical analysis was conducted using One-way ANOVA Tukey’s test,**** p < 0.0001. Control: 24-26 hours n=25, and 26-30 hours n=15. Irradiated: 24-26 hours n=11, 26-28 hours n =11 and 28-30 hours n=7 hours.

### Genetic ablation expands the period of damage-induced proliferative response during pupal development

Our results indicate that the cells in the pupal wing exhibit insensitivity to IR-induced apoptosis after 26 hours (APF). Consequently, it is plausible to consider that the absence of observed proliferative responses in irradiated pupae during the later stages of pupal development may be attributed to the ineffectiveness of IR in inducing apoptosis. To test this, we employed an alternative approach to effectively induce apoptosis. To this end we used the Gal4/UAS binary system with Gal80^ts^ to induce genetic ablation at different times APF. We overexpressed the pro-apoptotic gene *reaper* (*rpr*) in the posterior compartment of the pupal wing using the *hedgehog* (*hh*)*-Gal4* line. To ensure accurate and reliable results, we modified our protocols to accommodate differences in *Drosophila* development at different temperatures (see methods, Fig. 3A). We selected pupae at 0-4 hours APF, raised them at 17°C for varying lengths of time before shifting them to 29°C to activate *rpr* expression for a 20-hour period. To evaluate the proliferative response, we quantified the number of mitotic cells in pupal wings at 20-24 hours, 25-29 hours, 30-34 hours, and 35-40 hours APF, equivalent to a temperature of 25°C (Fig. 3A and methods).

**Figure 3.**
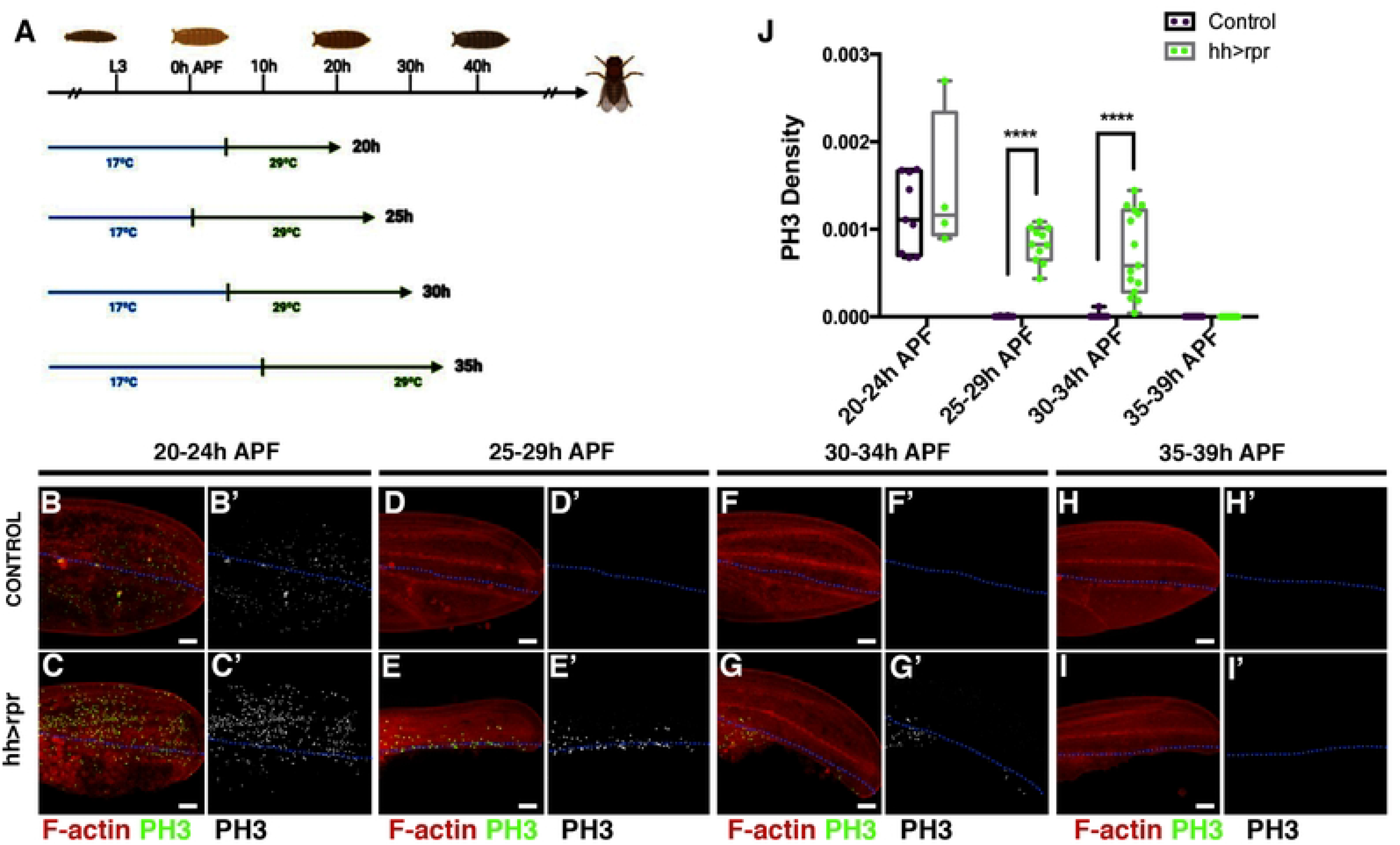
**Genetic ablation prolongs the proliferative period of pupal wing**. (A) Schematic diagram of the temperature shifts used in this experiment. (B-B’, D-D’, F-F’ and G-G’). Control pupal wing at 20-24 hours (B-B’), 25-29 hours (D-D’), 30-34 hours (F-F’) and 35-39 hours APF (H-H’). (C-C’, E-E’, G-G’ and I-I’) Ablated *UAS-rpr; hh-Gal4; tub-Gal80^ts^* pupal wing at 20-24hours (C-C’), 25-29 hours (E-E’), at 30-34 hours (G-G’) and 35-39 hours APF (I-I’). (J) The graph shows the density of mitotic cells (PH3 positive) in control wings and in damaged wings at different times APF. Error bars represent SEM. The discs were stained with anti-PH3 antibody (in green B-I and grey B’-I’) and Phalloidin (in red B-I). Statistical analysis was conducted using multiple comparation t-Test student (Mann-Whitney test) **** p<0.0001. Control: 20-24 hours n=9, 25-29 hours n=14. 30-34 hours n=10 and 35-39 n=10 hours. Ablated *UAS-rpr; hh-Gal4; tub-Gal80^ts^*: 20-24 hours n=4, 25-29 hours n=11. 30-34 hours n=15 and 35-39 n=11 hours. White scale bar 50μm.

Our results showed that, at 20-24 hours APF, damaged pupal wings exhibited an increase in cell proliferation compared to control wings, with most dividing cells located near the damaged area (Fig. 3 B-C’ and J). At 25-29 hours APF, no mitotic cells appeared in control wings, but we still observed a high number of mitotic cells in damaged wings (Fig. 3D-E’ and J). Similarly, at 30-34 hours, mitotic cells were present in damaged wings but not in control wings (Fig. 3F-G’ and J). However, we did not observe any dividing cells in either control or damaged wings after 35-39 hours APF (Fig. 3HI-I’ and J).

Interestingly, the overexpression of *rpr* during late-stage of pupal development resulted in the partial elimination of the posterior compartment, indicating that apoptosis was effectively induced at all experimental time points analyzed (Fig. 3). Consequently, the absence of a proliferative response after 35 hours (APF) could not be attributed to the lack of apoptosis. Therefore, our findings suggest that apoptosis triggers a proliferative response that lasts up to 30-34 hours APF, after which it declines. The mitotic signals produced in the damaged area may extend the proliferative period of pupal wing cells, but only until 30-34 hours (APF).

In summary, our study provides evidence that the duration of the proliferative response in pupal wing cells is limited and can be influenced by proliferative signals produce by dead cells.

### JNK signalling is activated in response to damage in late stages of pupal development

The JNK pathway plays a crucial role in triggering apoptosis-induced proliferative response during the larval stages (Bosch et al., 2005; Mattila et al., 2005; Ramet et al., 2002). To investigate the evolution of JNK pathway activity in response to cell death induction after metamorphosis, we examined the expression of the TRE-GFP reporter (Chatterjee and Bohmann, 2012) following the over-expression of *rpr* under the control of *hh-Gal4* at different times APF. We selected pupae at 0 hours APF and raised them at 17°C until they reached the equivalent of 10-15 h APF at 25°C. Next, we induced ectopic expression of *rpr* by transferring them to 29°C until they reached the equivalent of 25-30 h, 30-35h, and 35-40 h APF at 25°C (see Fig. 4A). At 25-30 h and 30-35 h APF, TRE-GFP expression was observed in the damaged region, unlike the controls where the expression of this reporter was restricted to hemocytes located in the wing veins (see Fig. 4B-E’). Interestingly, at 35-40 h AFP, the expression of the TRE-GFP reporter persisted in the damaged region even though cell proliferation was no longer activated at this age by cell death induction (see Fig. 4F-G’).

**Figure 4.**
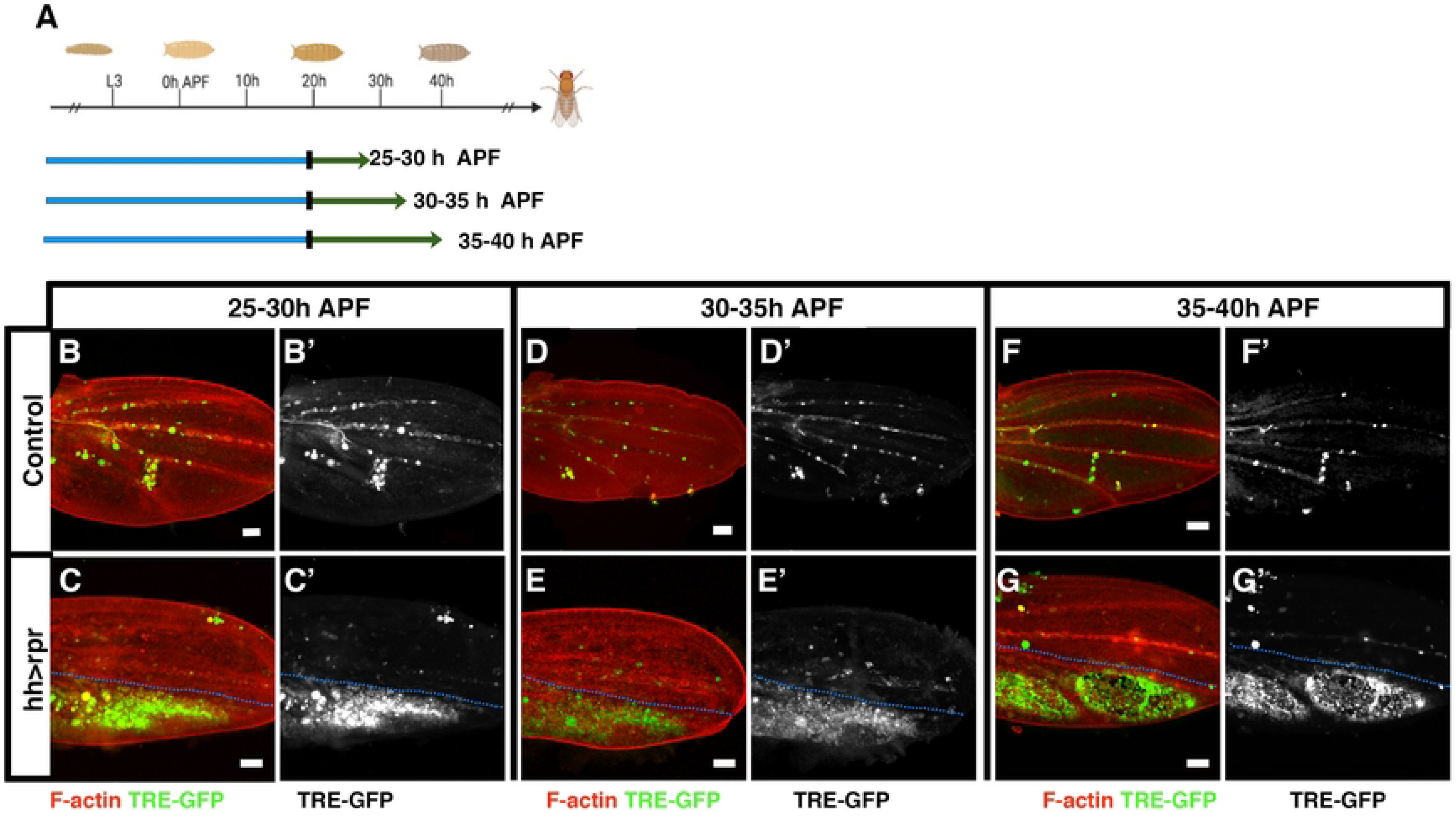
**Pattern of JNK activation following genetic ablation of the posterior compartment in the wing disc**. (A) Schematic diagram of the temperature shifts used in this experiment. (B-B’, D-D’, and F-F’). *hh-Gal4; tub-Gal80^ts^* TRE-GFP control pupal wings at 25- 30 hours (B-B’), 30-35 hours (D-D’), and 35-40 hours APF (F-F’). (C-C’, E-E’, and G-G’) Ablated *UAS-rpr; hh-Gal4; tub-Gal80^ts^* TRE-GFP pupal wings at 25- 30 hours (C-C’), 30-35 hours (E-E’), and 35-40 hours APF (G-G’). Note that while in control discs the expression of the JNK pathway reporter is restricted to hemocytes, in the genetically ablated discs, the JNK pathway reporter is expressed in the apoptotic cells of the posterior compartment. The expression of the reported is shown in green. The blue dotted line indicates the boundary between the anterior and posterior compartment. White scale bar 50μm

Our experiments using IR to induce apoptosis (Fig. S2) yielded similar results. We observed JNK activation in response to irradiation, regardless of whether it occurred during early stages, when cell proliferation is activated, or late stages, when it is not. These findings suggest that the loss of proliferative response during late stages of pupal development is not caused by an inability of cells to activate JNK. To test this idea, we overexpressed a constitutively activated form of the JNK-kinase Hemipterous (*hep^CA^*) in late stages of pupal development. Our results showed that *hep^CA^*overexpression induced a proliferative response only until 30-35 hours AFP, and not at later stages (Fig. 5).

**Figure 5.**
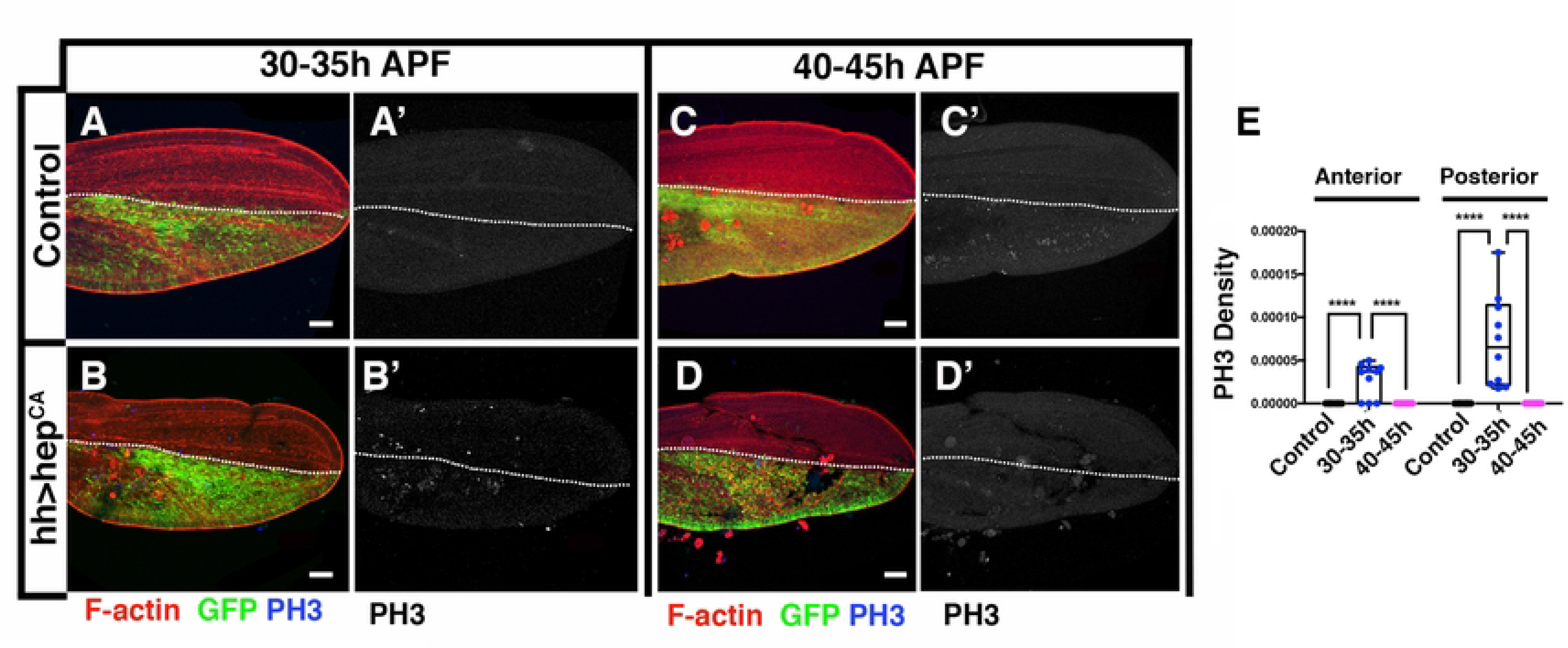
The ectopic activation of *hep^CA^* is not sufficient to expand the proliferative period of pupal wing beyond 30-35hous APF. (A-A’, C-C’) *hh-Gal4; tub-Gal80^ts^* control pupal wings at 30-35 hours (A-A’), and 35-40 hours APF (C-C’). (B-B’, C-C’) *UAS-hep^CA^; hh-Gal4 UAS-GFP; tub-Gal80^ts^* pupal wings at 30-35 hours (B-B’), and 35-40 hours (C-C’). (E) The graph shows the density of mitotic cells (PH3 positive) in control wings and in *hh-Gal4 UAS-GFP; tub-Gal80^ts^* wings at different times APF. Error bars represent SEM. The discs were stained with anti-PH3 antibody (in blue A-D, and grey A’-D’) and Phalloidin to reveal F-Actin (in red B-I). Statistical analysis was conducted using multiple comparation t-Test student (Mann-Whitney test) **** p<0.0001. Anterior compartment: Control n=10, and *UAS-hep^CA^; hh-UAS-GFP; tub-Gal80^ts^*30-35 hours n=8, and 35-40 hours n=10. Posterior compartment: Control n= 10, and *UAS-hep^CA^; hh-Gal4 UAS-GFP; tub-Gal80^ts^* 30-35 hours n=8, and 35-40 hours n=10. The white dotted line indicates the boundary between the anterior and posterior compartment. White scale bar 50μm

### The over-expression of *CycE* or *E2F1* in damaged pupal wing is not sufficient to promote a proliferative response after 35-40 hrs APF

Previous studies have shown that the overexpression at pupal stages of key regulators of the G1-S transition, such as CycE, CycE/Cdk2, CycD/Cdk4 or E2F1 maintains cell division until 30 h APF, but not later (Buttitta et al., 2007). This is due to a positive feedback mechanism between CycE and E2F1 that is active during larval and early pupal development, but ceases after 30 hrs APF (Buttitta et al., 2007). Similarly, our observations indicate that apoptosis can trigger proliferation up to 30-35 hours AFP, but not beyond. Thus, it is plausible that the mitogenic signals generated by apoptotic cells specifically activate CycE/Cdk2 or E2F1, thereby triggering a proliferative response that is sustained only until approximately 30-35 hours APF. This time frame coincides with the period when the feedback mechanism between CycE and E2F1 is still functioning. To test this hypothesis, we utilized the double transcriptional trans-activator system *sal^E/Pv^-LHG/lexOp* in combination with *Gal4/UAS* to overexpress *E2F1 dp* or *CycE* while inducing apoptosis. The *sal^E/Pv^-LHG* transgene contains a *Gal80* suppressible form of *LexA* (Yagi et al., 2010). To conditionally express both the *sal^E/Pv^-LHG/lexOp* and *Gal4/UAS* binary systems, we used *tub-Gal80^ts^*. We induced apoptosis in the *sal^E/Pv^-LHG* domain of the wing disc by using *lexOp-rpr*, while simultaneously overexpressing *UAS-CycE* or *UAS-E2F1* in the overlapping anterior compartment with the *cubitus interruptus* (*ci*)-*Gal4* line (Fig. 6A).

**Figure 6.**
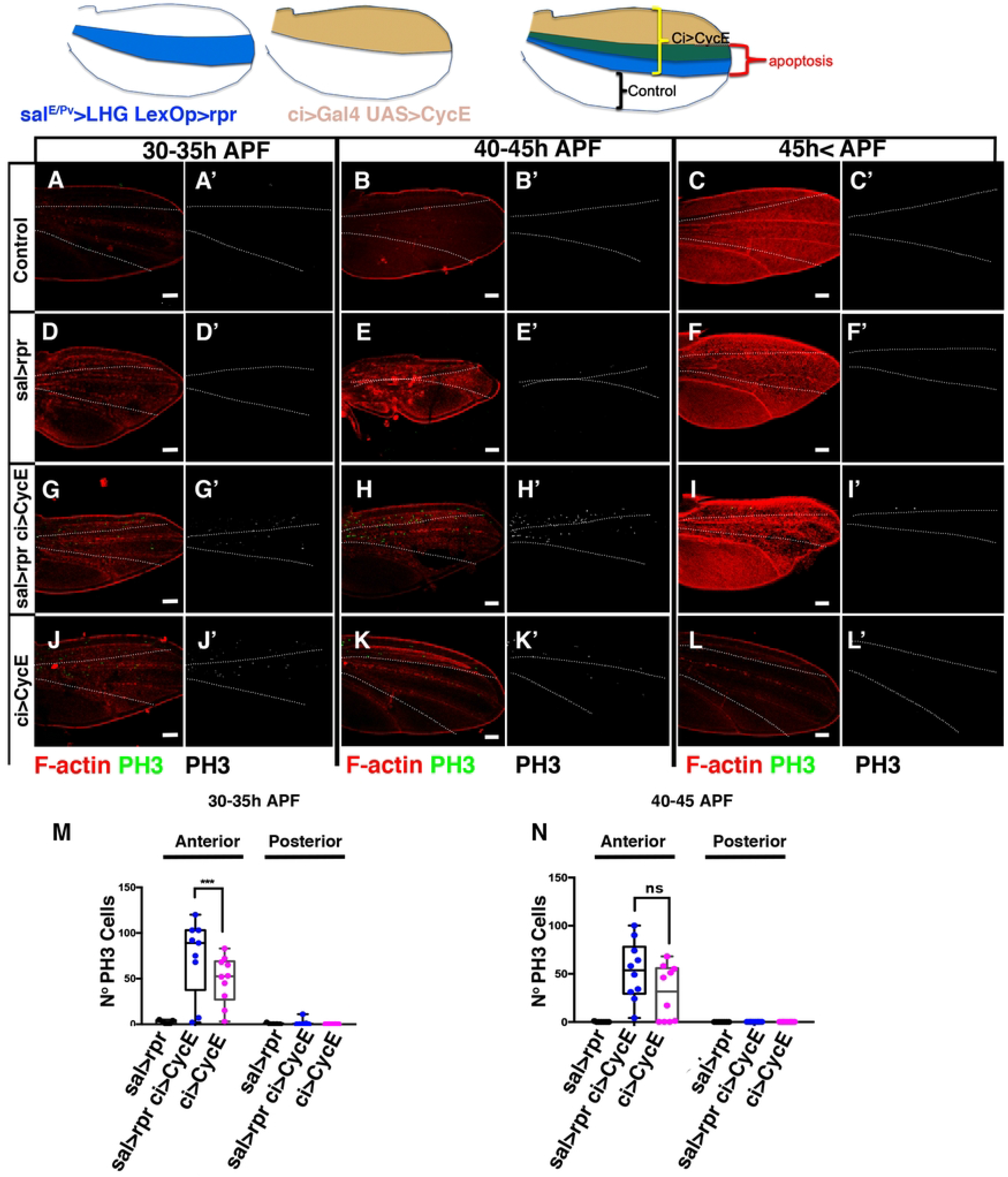
CycE is not a limiting factor for apoptosis-induced proliferation during pupal development. (A-C’) Control pupal wings at 30-35 hours (A-A’), 40-45 hours (B-B’) and ≥45 hrs APF (C-C’). (D-F’) *lexOp-rpr; ci-Gal4; sal^E/Pv^-LHG tub-Gal80^ts^* pupal wings at 30-35 hours (D-D’), 40-45 hours (E-E’) and ≥45 hours APF (F-F’). (G-I’) *lexOp-rpr; ci-Gal4 ;sal^E/Pv^-LHG tub-Gal80^ts^ UAS-CycE* pupal wings at 30-35 hours (G-G’), 40-45 hours (H-H’) and <u>> 45hrs APF</u> 45 hrs< APF (I-I’). (J-L’) *ci-Gal4 tub-Gal80^ts^ UAS-CycE* pupal wings at 30-35 hours (J-J’), 40-45 hours (K-K’) and 45 hrs< APF (L-L’). (M-N) The graphs show number mitotic cells (PH3 positive) in the different experimental condition analysed. The discs were stained with anti-PH3 antibody (in Green A-L, and grey A’-L’) and Phalloidin to reveal F-Actin (in red B-I). Statistical analysis was conducted using multiple comparation t-Test student (Mann-Whitney test) *** p<0.001. (M) 30-35h Anterior compartment: *sal>rpr* n=5, *sal>rpr ci>CycE* n=9, *ci>CycE* n=10. Posterior compartment *sal>rpr* n=5, *sal>rpr ci>CycE* n=9, *ci>CycE* n=10. (N) 40-45h Anterior compartment: *sal>rpr* n=7, *sal>rpr ci>CycE* n=10, *ci>CycE* n=10. Posterior compartment *sal>rpr* n=7, *sal>rpr ci>CycE* n=10, *ci>CycE* n=10 The expression domain of *sal* is outlined by the white dotted line. White scale bar 50μm

We selected pupae between 0-5 hours old APF and maintained them at 17°C until they reached the developmental stage equivalent to 25-30 hours APF at 25°C. Following this, we shift them to 29°C to trigger ectopic expression of *UAS-CycE* (*ci-Gal4, UAS-CycE*), while simultaneously inducing apoptosis (*sal^E/Pv^-LHG/lexOp-rpr)*. The discs were analysed at the developmental stage equivalent to 35-40 hours APF or beyond 40 hours APF. Our observations revealed a high number of mitotic cells in the anterior compartment of discs expressing *CycE*, or co-expressing *rpr* and *CycE*, at the developmental stages equivalent to 35-40 hours and 40-45 hours AFP (as illustrated in Fig. 6). However, at 45 hours APF, we did not observe any mitotic cells in either the *CycE*-expressing discs or the discs co-expressing *CycE* and *rpr*. We obtained similar results when we induced apoptosis and overexpressed *E2F1* simultaneously (Fig. S4).

All together these results suggest that damage can only induce a proliferative response while the positive feedback mechanism between CycE and E2F1 operates.

### Ecdysone-responsive transcription factor E93 blocks the proliferative response during pupal development

The end of the positive feedback loop between CycE and E2F, as well as the ending of the proliferative response following damage, coincide with the initiation of epigenetic silencing of the regulatory regions of critical genes involved in cell cycle regulation, including *CycE*, *E2F1*, and *string* (*stg*) (Uyehara et al., 2017; Guo et al., 2016; Ma, 2019). In *Drosophila*, developmental transitions are regulated by the hormone ecdysone, and ecdysone-responsive transcription factors control temporal changes in chromatin accessibility during wing disc development (Uyehara et al., 2017; Guo et al., 2016; Ma, 2019). Specifically, the E93 transcription factor is transcriptionally activated at 18 hours and 24 hours APF (Uyehara et al., 2017). Loss-of-function mutations in the *E93* gene results in chromatin accessibility changes at several genome regions in 24 h and 44 h APF wing discs (Uyehara et al., 2017). Importantly, the progressively closed chromatin status observed at regulatory regions of the *CycE*, *E2F1* and *stg* genes between third instar and 44 hours APF is attenuated in *E93* mutants (Uyehara et al., 2017) (Fig S5). Given that this temporal window coincides with the end of apoptosis-induced proliferation during pupal development, it is plausible that E93, by blocking the chromatin accessibility at these cell cycle regulators, prevents the induction of this response. To explore this idea, we have irradiated *E93* mutant larvae at 20 hours APF and examined the proliferative response 20 hours later (40 hours APF). However, in non-irradiated E93 mutant pupal wings, we still observed some PH3 positive cells, confirming the role of E93 in suppressing the proliferative capability of wing disc cells at this specific time point (Fig. 7A and B). This finding highlights the importance of E93 in regulating cell proliferation during pupal wing development (Fig. 7A and B). Our results strongly suggest that E93, through its involvement in the ecdysone response, acts as a key orchestrator of the intricate cellular processes that regulate and limit the proliferative response of pupal cells towards the end of their development.

**Figure 7.**
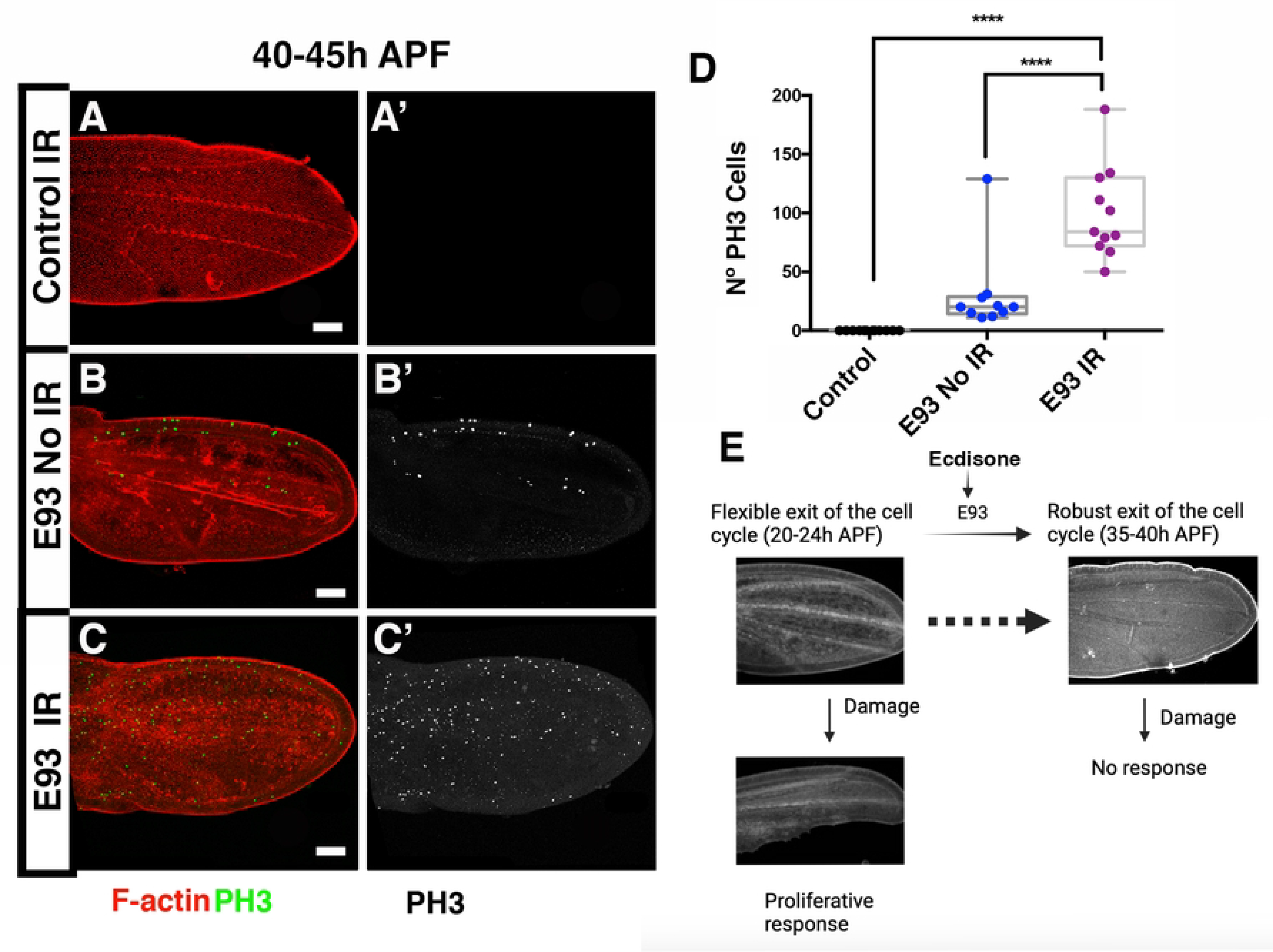
In *E93* mutant damage prolongs proliferative response beyond 40 h APF. (A-C’) Pupal wings at 40-45 hours APF. Control (A-A’), *E93* non-irradiated pupal wings (B-B’) and *E93* irradiated pupal wings (C-C’). (D) The graphs show number mitotic cells (PH3 positive) in the different experimental condition analysed. The discs were stained with anti-PH3 antibody (in Green A-C, and grey A’-C’) and Phalloidin to reveal F-Actin (in red A-C). (E) A proposed model for the dynamic response to damage during pupal development. Statistical analysis was conducted using One-way ANOVA Tukey’s test,**** p<0.0001. Control n=12, *E93* non-irradiated n= 10, *E93* irradiated n= 11. White scale bar 50μm

## Discussion

Regeneration is a complex process that relies on multiple cellular and molecular mechanisms working in concert to uphold tissue homeostasis and effectively respond to external challenges. One critical aspect of this process is the induction of regenerative growth, which is often lost during development or aging. In *Drosophila* development, multiple studies have shown that cell proliferation is one of the primary responses during discs regeneration in larval stages (Adler and Macqueen, 1984; Dale and Bownes, 1981; Kiehle and Schubiger, 1985; Bryant and Fraser, 1988; Sustar and Schubiger, 2005; Bosch et al., 2008; Worley and Hariharan, 2022). When damage occurs, apoptotic cells generated by the insult can initiate a process known as Apoptosis-induced proliferation (Perez-Garijo and Steller, 2015; Ryoo HD, 2012). Different studies using third instar larval imaginal discs have suggested that this ability is lost at the end of larval stage or during pupal development (Smith-Bolton et al., 2009; Diaz-Garcia and Baonza, 2013). However, our work shows that damage induction in the wing disc produces a proliferative response that is maintained up to 30 hours APF.

Previous studies have demonstrated that overexpression of key regulators of the G1-S transition, such as CycE, CycE/Cdk2, CycD/Cdk4, or E2F1, in pupal stages can maintain cell division until 30 hours APF. However, overexpression of these regulators after that time is not sufficient to activate cell proliferation. To keep cells proliferating until at least 40-44 hours APF, ectopic activation of both CycE and E2F1 is necessary. It has been proposed that this is due to the existence of a positive feedback mechanism between CycE and E2F that is active during larval and early pupal development, but finishes after 30 hours APF (Buttitta et al., 2007; Guo et al., 2016; Ma, 2019). Our findings reveal that induction of apoptosis in wing imaginal discs before 24 hours APF induces cell proliferation up to 30 hours APF, which coincides with the end of the positive feedback between CycE and E2F1. This suggests that mitogenic signals released by apoptotic cells would specifically act on CycE or E2F1, stimulating cell division during the active positive feedback mechanism. However, these signals alone are insufficient to counteract the signals that drive cell cycle exit at the conclusion of the proliferative phase.

### Hormones block apoptosis-induce proliferation

In *Drosophila* developmental transitions are regulated by the hormone Ecdysone. The signals induce by this hormone are mediated by the Ecdysone-responsive transcription factors, that are involved in regulating temporal changes in chromatin accessibility that occur throughout wing disc development (Uyehara et al., 2017; Guo et al., 2016; Ma, 2019). Specifically, the E93 transcription factor is expressed at 18 hours and 24 hours PFD APF (Uyehara et al., 2017). Mutation of the *E93* gene results, not only in the loss of accessibility in several regions of the genome, but also in the aberrant presence of numerous regions of open chromatin at 24 and 44 hours APF wing discs including regulatory regions adjacent to the cell cycle regulator *CycE*, *E2F1* and *stg*. This suggests that the elevated levels of the hormone ecdysone during pupal development trigger the activation of E93.Consequently, E93 induces the irreversible exit of wing disc cells from the cell cycle by epigenetically silencing the regulatory regions of crucial cell cycle regulator genes.

During pupal development, the induction of cell death triggers the activation of JNK pathway, both in early (Bosch et al., 2005) and late stages, even when apoptosis-induced proliferation is not observed. Although the JNK pathway is known to suppress the Polycomb group of proteins (Lee et al., 2005), our results suggest that its activation in pupal discs is not enough to overcome epigenetic silencing of key cell cycle regulators.

Our data provide evidence supporting a model in which ecdysone-mediated developmental programming induces specific alterations in chromatin accessibility of crucial cell cycle regulator genes (Uyehara et al., 2017; Guo et al., 2016; Ma, 2019). These alterations, in turn, govern the process of cell cycle exit and subsequently suppress the proliferative response that is normally activated in response to damage (Fig 7 E). This model is likely to be applicable to other organisms, such as urodeles, zebrafish, and mice. For instance, Lin-28, an RNA-binding protein that regulates Let-7, has been shown to inhibit the expression of thyroid hormone target genes, delay development, and prolong regenerative potential after damage in urodeles (Faunes et al., 2017). Similarly, inhibiting thyroid hormone signaling at the level of its synthesis and at the receptor level in neonatal mice rescues the proliferative capacity of cardiomyocytes and the regenerative potential of the adult heart (Hirose et al., 2019). Conversely, exogenous administration of thyroid hormones in zebrafish inhibits the regenerative capacity of the heart and caudal fin (Sun et al., 2019). The result presented in this work highlights the intricate interplay between developmental cues, chromatin modifications, and the regulation of cell cycle dynamics, shedding light on the mechanisms underlying the fine-tuned control of cellular proliferation and regeneration

## Materials and Methods

### Fly strain

The following strains were used in this study. Unless otherwise was indicated, strain descriptions can be found at http://flybase.bio.indiana.edu.

*w; en-Gal4 UAS-GFP; tub-Gal80^ts^, w; tub-Gal80^ts^; hh-Gal4, w; ci-Gal4; sal^E/Pv^ LHG, tub-Gal80^ts^ (Santabarbara-Ruiz et al., 2015), UAS-CycE(II), UAS-E2F1, UAS-Dp, UAS-CycE lexOp-rpr, UAS-E2F1 UAS-dp lexOp-rpr, lexOp-rpr* (Santabarbara-Ruiz et al., 2015)*, TRE-DsRed, UAS-hep^CA^*

### Protocol for Genetic ablation and irradiation experiments

The development of pupae is observed to be 2.5 times slower at 17˚C compared to 25˚C and 1.3 times faster at 29˚C compared to 25˚C, as per our experimental conditions. To ensure equivalent stages of development, all time points for pupae grown at 17˚C were adjusted to match the stages at 25˚C.

Figure 1: White prepupae were collected over a 4-hour period for staging. They were irradiated at 0-4, 10-14, and 20-24 hours after pupa formation (APF) and analyzed 20 hours later. Throughout pupal development, they were kept at a constant temperature of 25˚C.

Figure 2: Pupae were selected at 2-hour intervals, starting at 20 hours APF. The selected pupae were immediately irradiated, and their apoptotic response was analyzed 4 hours later.

Figure 3: White prepupae were collected at 4-hour intervals for staging and kept at 17˚C for different durations. Subsequently, the pupae were shifted to 29˚C until reaching the equivalent developmental time points of 20-24 hrs, 25-29 hrs, 30-34 hrs, or 35-39 hrs APF at 25˚C for dissection.

Pupae 20 -24--- (10hrs at 17°C) + (12hrs at 29°C)

Pupae 25 -29--- (20hrs at 29°C)

Pupae 30-34 --- (10hrs at 17°C) + (20hrs at 29°C)

Pupae 35-39 --- (20hrs at 17°C) + (20hrs at 29°C)

Figure 4.-White prepupae were collected at 5-hour intervals for staging and kept at 17˚C for different durations. Subsequently, the pupae were shifted to 29˚C until reaching the equivalent developmental time points of, 25-30 hrs, 30-35 hrs, or 35-40 hrs APF at 25˚C for dissection.

Pupae 25 -30--- (20hrs at 17°C) + (12hrs at 29°C)

Pupae 30-35 --- (20hrs at 17°C) + (16hrs at 29°C)

Pupae 35-40 --- (20hrs at 17°C) + (20hrs at 29°C)

Figure 5.- White prepupae were collected at 5-hour intervals for staging. The temperature shifts used in this experiment was:

Pupae 30-35 --- (30hrs at 17°C) + (16hrs at 29°C)

Pupae 40-45 --- (48hrs at 17°C) + (16hrs at 29°C)

Figure 6.-White prepupae were collected at 5-hour intervals for staging. The temperature shifts used in this experiment was:

Pupae 30-35 --- (30hrs at 17°C) + (16hrs at 29°C)

Pupae 40-45 --- (48hrs at 17°C) + (16hrs at 29°C)

Pupae 45< (52hrs at 17°C) + (21hrs at 29°C)

Figure 7.-Pupae 40-45 :Throughout pupal development, they were kept at a constant temperature of 25˚C.

### Irradiation

Pupae were given a dose of 4000 R using Philips-MG-102 irradiation unit.

### Immunocytochemistry

Immunostaining of the wing discs was performed according to standard protocols. The following primary antibodies were used: rabbit anti-Phospho-histone 3 1:200 (Cell Signalling Technology), rabbit anti-cleaved Dcp1 1:200 (Cell Signalling Technology), Phalloidin-TRITC 1:200. They were obtained from the Developmental Studies Hybridoma Bank at the University of Iowa. Secondary antibodies were used at dilutions of 1:200.

Imaginal discs were mounted in Vectashield mounting fluorescent medium (Vector Laboratories, Inc.).

## Acknowledgements

We thank Jose Felix de Celis, and Luis Alberto Baena for providing reagents and useful discussion. We are very grateful to Florenci Serras, the Bloomington Stock Center and the Developmental Studies Hybridoma Bank for providing fly strains and antibodies. This study was supported by a grant from: FEDER/ Ministerio Ciencia-Agencia Estatal de Investigación PID2021-127114NB-I00 and institutional grant from Banco de Santander to the CBMSO.

